# Replay without sharp wave ripples in a spatial memory task

**DOI:** 10.1101/2024.05.13.593931

**Authors:** John Widloski, David J. Foster

## Abstract

Sharp-wave ripples in the hippocampus are believed to be a fundamental mechanism for the consolidation of episodic memories. During ripples, hippocampal neurons are re-activated in sequences called replay, which have been hypothesized to reflect episodic memory content. Ripples and replay are usually reported to co-occur, and are commonly thought to reflect the same process. Here we report that, in rats performing an open field spatial memory task, replays readily occur in the complete absence of ripples. Moreover, the occurrence of ripple-less and ripple-containing replays is not random, but precisely organized in terms of virtual space: Ripples are confined to “ripple fields”, which are spatially-restricted areas defined over the virtual locations depicted during replay and independent of the actual location of the animal. Similar to allocentric coding by place fields, ripple fields are independent of the direction of travel, and stable throughout the recording session. Ripple fields track changes to environmental structure caused by the addition or subtraction of barriers to movement, consistent with ripples conveying information about the incorporation of novel experiences. Moreover, ripple fields were matched across different rats experiencing the same barrier configuration, highlighting the robustness of the ripple field spatial code. We hypothesize a new relationship between ripples and replay, in which a subset of replays that is particularly relevant to learning or novelty is paired with ripples, in order to promote its selective broadcast to the rest of the brain for consolidation.

## INTRODUCTION

Sharp wave ripples are among the most salient events in the mammalian brain, constituting a transient (∼100 ms) high frequency (100-200 Hz) ripple in the local field potential reflecting a highly coordinated, synchronous firing of neurons across brain regions (Chrobak & Buzsáki, 1994; Qin et al., 1997; Pennartz et al., 2004; Lansink et al., 2009; Peyrache et al., 2009; Girardeau 2017; Sjulson et al., 2018) that is well suited for the induction of synaptic plasticity (Buzsáki, 1986; Bi & Poo, 1998; Debanne et al., 1998; Wittenberg, 2006). Because ripples occur primarily during rest/sleep and tend to start in the hippocampus, an area long associated with memory (Scoville & Milner, 1957) and spatial mapping (O’Keefe & Nadel, 1977), ripples are thought to play a crucial role in episodic memory, by binding complex sensory-motor memory traces to place information in the hippocampus for the purposes of memory recall and consolidation (Csicsvari & Dupret, 2014; Buzsáki, 1989; Sutherland & McNaughton, 2000; Squire et al., 2015; Teyler & Discenna, 1986). At the same time, hippocampal activity during ripples is highly structured, encoding continuous paths through the environment that evoke the animals own movement through space, albeit at speeds 20x faster (Foster & Wilson, 2006; Diba & Buzsáki, 2007; Pfeiffer & Foster, 2013; Davidson et al., 2009; Widloski & Foster, 2022). These two phenomena, ripples and replay, are believed to function hand in hand during episodic memory encoding, the former broadcasting the memory contents of the latter, as well as facilitating its consolidation. Supporting this hypothesis, ripples are known to facilitate long-term potentiation and reorganization of hippocampal synapses (Sadowski et al., 2016; Rolotti et al., 2022) and blocking replay-associated sharp wave ripples has also been shown to lead to deficits in both memory consolidation and retrieval (Jadhav et al., 2012; Singer et al., 2013; Girardeau & Zugaro, 2009; Maingret et al., 2016; Ego-Stengel & Wilson, 2010).

However, the precise nature of the relationship between ripples and replay remains elusive. For example, replay detectors are generally predicated on the existence of ripples or bursts (i.e., supra-threshold behavior in the ripple power or population spike density; O’Neill et al., 2017; Pfeiffer & Foster, 2013; Denovellis et al., 2021; Maboudi et al., 2018; Stella et al., 2019; Tingley & Peyrache, 2020). However, it is possible that replays may exist covertly but have gone undetected. Indeed, compressed hippocampal sequences in the absence of ripples or bursts are readily observable during run (Dragoi & Buzsáki, 2006; Foster & Wilson, 2007; Gupta et al., 2012; Wikenheiser & Redish, 2015), though shorter in length than replay, suggesting that ripples are not required for hippocampal sequence generation. In addition, ripples and replay are not always one-to-one: When animals are exposed to long linear tracks or complex environments, replays can be very long, in fact much longer than the timescale of a single ripple (Davidson et al., 2009; Widloski & Foster, 2022). In these cases, multiple ripples tend to occur (Buzsáki et al., 1983; Suzuki & Smith, 1987; Davidson et al., 2009; Yamamoto & Tonegawa, 2017; Wu et al., 2014), suggesting that ripples may be building blocks of replay. Taken together, these considerations suggest that the relationship between replay and ripples may be complex.

To address these questions, we developed a ripple-independent replay detector that enabled us to track ripples and replay independently and applied it to a large data set consisting of hundreds of simultaneously recorded place cells from rats performing a spatial memory task. Crucially, the task was designed to elicit extended replay in two spatial dimensions, and perturb it with repeated manipulations of spatial and reward contingencies, to generate a wide variety of replay patterns. We find that the hippocampus participates in the encoding of long spatial sequences in the complete absence of ripples. We also show that ripple timing is tightly constrained by the content of replay, occurring in spatially restricted “ripple fields” as a function of the encoded location during replay, and context dependent, with ripple fields “remapping” across environments when the barriers were moved. Our results demonstrate a finely tuned hierarchical relationship between ripples and replay whereby replay content regulates the occurrence of ripples, while replays can occur independently of ripples.

## RESULTS

### Replays can occur in the complete absence of ripples

To determine whether replay content can exist in the absence of bursts or ripples, we utilized a previously published data set (Widloski and Foster, 2022) comprising of 7273 replays recorded across 37 session from 3 rats and applied a novel replay detection method that did not depend on the occurrence of population bursts or ripples. Briefly, during each session, rats were trained on a spatial memory task to search for liquid chocolate in a square arena with movable barriers. Chocolate was available in one of 9 food wells, which alternated on consecutive trials between a learnable fixed location (Home well) and other unpredictable locations (Random wells). We recorded the activity of up to 295 hippocampal place cells simultaneously (mean per session = 182 cells). Place fields were determined for each active cell by normalizing the spatial distribution of rat positions at spike times during run by the rat position occupancy, and memory-less Bayesian position estimation was used to decode the posterior probability of position from the spiking of all simultaneously recorded cells. During stopping periods in the task, candidate events were identified as continuous epochs lasting at least 100 msec where the decoded position changed smoothly. Those candidate events that satisfied spatial coverage criteria were classified as replays, but otherwise no requirements related to sharp-wave ripple power or population spiking were imposed.

Surprisingly, replays possessing long-duration, smooth, continuous spatial content were readily detected that were not accompanied by population bursts or ripples (**Figure 1A,C**). Ripple power was defined as the average ripple power across all tetrodes with at least 4 active place cells (i.e., were close to the hippocampal cell layer). For these ripple-less events, peak ripple power within the event was no different from a shuffle distribution derived from the peak ripple power from 100 random equal-length snippets of LFP during stopping periods but outside of ripple times. Ripple-less replays were not due to poor tetrode placement or averaging ripple power across tetrodes, as replays with strong population bursts and coincident across-tetrode ripples were readily detected (**Figure 1B,D**). Indeed, replays with vs. without ripples could occur only seconds apart (**Figure S1**).

**Figure 1.**
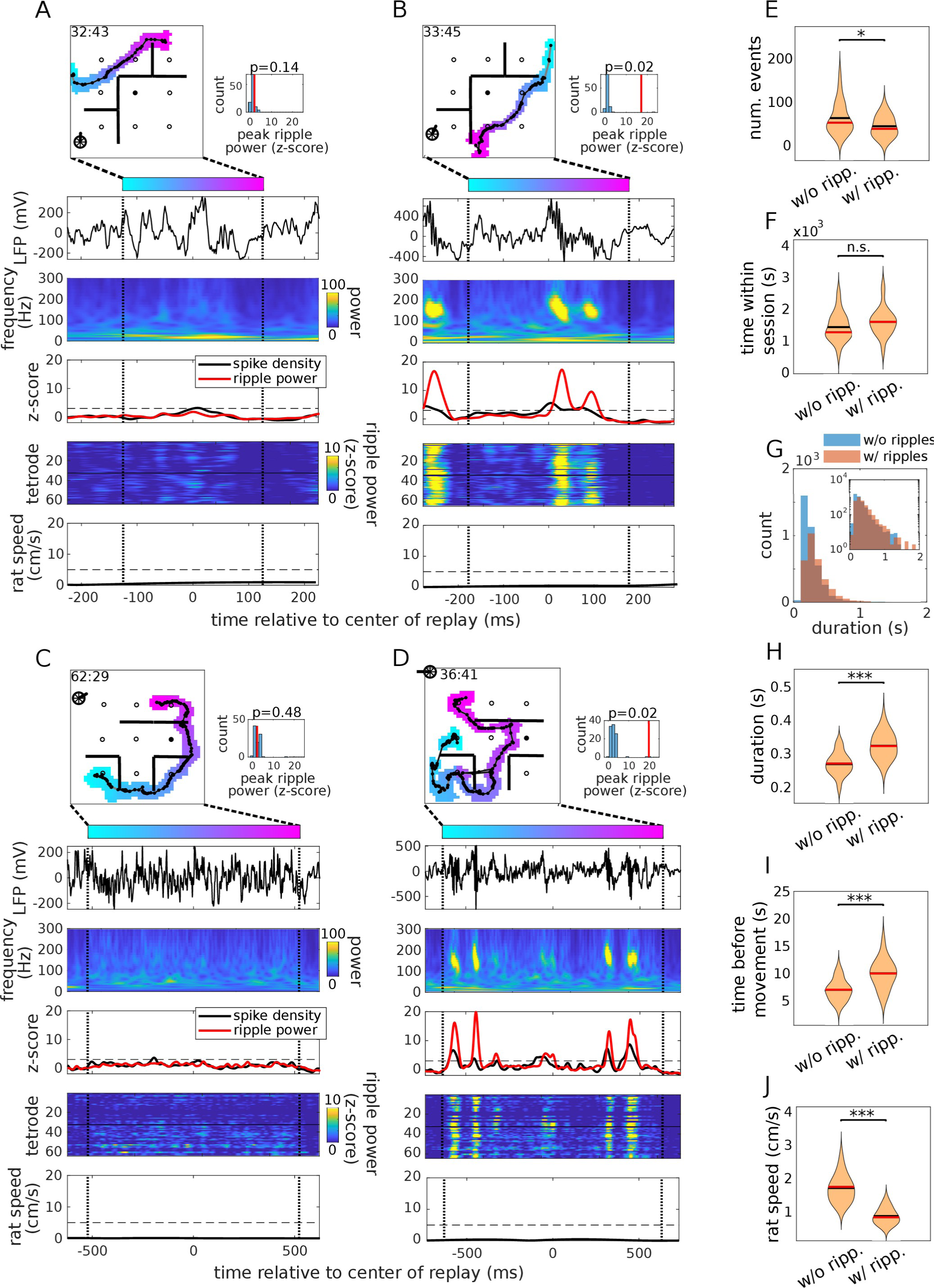
Replays can occur in the complete absence of ripples. (A) Top row: Example replay from rat 1, session 75. The colored blob is the posterior probability of a replay thresholded at 0.01 and color-coded according to elapsed time within the replay. Solid black line: replay center-of-mass. Time within session is shown at the upper left (min:sec). Black circles and straight lines are the locations of reward and transparent barriers, respectively. Filled circle is the home well. Right: Peak ripple power (vertical line) vs. shuffle, where shuffle is the peak ripple power (z-scored and averaged across tetrodes) from 100 random equal-length snippets of LFP during stopping periods but outside of ripple times, with p-value indicated above. Second row: Raw LFP trace from single electrode as a function of time within the replay. Black vertical dashed lines demarcate start and end of replay. Third row: Spike density summed across all recorded cells (black) and ripple power averaged across tetrodes (red). Dashed horizontal line indicates z-score of 3 and is standard detection criteria for ripple/spike density events. Fourth row: Ripple power across tetrodes. Tetrodes 1-32 (33-64) targeted the left (right) hemisphere. Fifth row: Rat speed. Dashed horizontal line indicates rat speed of 5 cm/s. (B) Replay with ripples, recorded in the same session as (A) a minute later during the same stopping period. (C-D) Replays without (C) and with (D) ripples from rat 2, session 87. (E) Total numbers of replay detected with vs. without ripples, averaged across sessions (n = 37 sessions). Likewise, (F) replay duration, (G) time of replay relative to rat movement (before or after), (H) time of replay within session, and (I) rat speed.

The number of ripple-less replays was comparable to the number of replays with ripples (**Figure 1E**), and were distributed in a likewise manner as a function of time within the session (**Figure 1F**) and duration, with long duration ripple-less replays occurring nearly as frequently as replays with ripples (**Figure 1G**). While ripple-less replays tended to be on average shorter in duration and occur closer to movement onset (**Figure 1G-H**), they nonetheless were associated with very low rat speeds (less than 1 cm/sec on average; **Figure 1I**) and had relatively long durations on average (∼270 msec), ruling out that these events are just theta sequences. In sum, we found a large number of replay events occurring during immobility that would have gone undetected if classical methods of replay detection had been used.

### Ripples occur reliably as a function of encoded location during replay

To determine whether there was any relationship between ripple timing and replay content, we plotted all replays from a given session with the ripple power for each replay superimposed (**Figure 2B**). Strikingly, ripples did not occur randomly as a function of replay location but were spatially restricted. We defined the session “ripple field” by computing, for each spatial bin in the environment, the sum total ripple power in that bin divided by the replay occupancy (**Figure 2C**). Ripple fields were found to exhibit spatial information and stability that was greater than chance for most sessions (spatial information: 30 out of 37 sessions; stability: 37 out of 37 sessions; **Figure 2D**), suggesting that ripple timing was controlled primarily by the position of replay in the environment. Here, shuffles were computed by circularly permuting the ripple power across all concatenated replays (**Figure S2**). When ripple times were temporally shifted, spatial information dropped precipitously within tens of milliseconds (**Figure 2E**), indicating that ripple timing during replay was relatively precise. Ripple fields did not simply reflect replay or rat location accumulation zones (0 out of 37 sessions and 4 out of 37 sessions respectively; **Figure 2I**), both of which tended to concentrate around reward locations (**Figure 2H**). We also checked whether ripple occurrence within the ripple field depended on the direction of replay travel. For each session, we extracted the location of each ripple field and compared the distribution of angles associated with replay entrance into the field with and without ripples. We found that for most sessions, these distributions were matched (**Figure 2I**), indicating that ripples tended to occur independently of the direction of replay travel.

**Figure 2.**
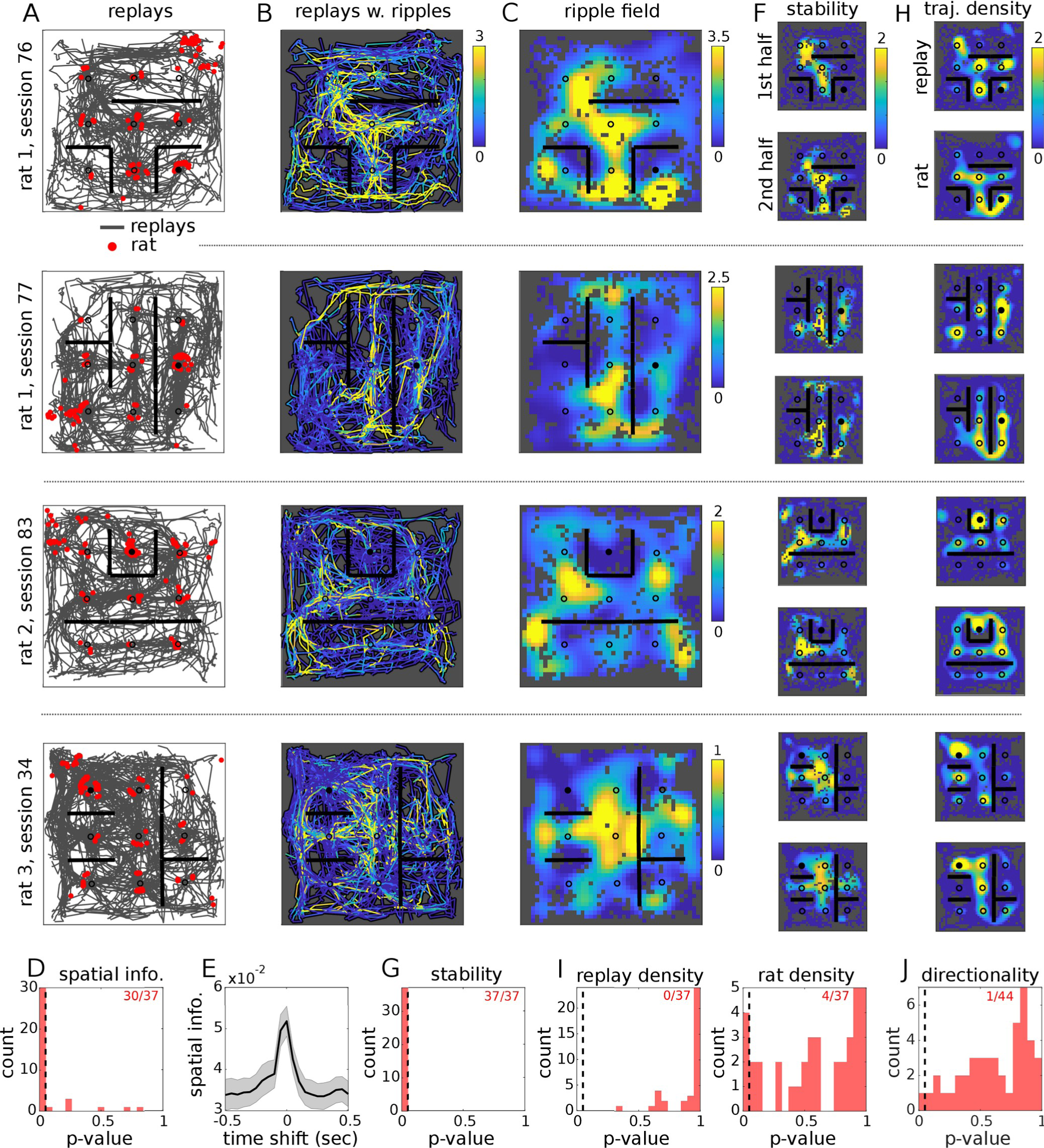
Ripples occur reliably as a function of encoded location during replay. (A) Replays from four example sessions across three rats. Each panel shows all replays recorded within the session (gray traces are replay center-of-masses). Red dots indicate rat location at time of each replay. (B) Replays in (A) with ripple power superimposed. Ripple power was computed by averaging across tetrodes. (C) Ripple fields, computed by averaging ripple power across spatial bins in (B) and normalizing by replay occupancy. Gray bins indicate locations with zero replay occupancy. (D) Significance of ripple field spatial information measured across sessions (n = 37 sessions). Null distribution computed on ripple fields after circularly permuting ripple power across replay events. Number of significant sessions indicated at upper right. (E) Ripple field spatial information as a function of temporally-shifted ripple power, averaged across sessions (n = 37 sessions). (F) Ripple fields derived from the first vs. the second half of replays (chronologically) in (B). Matrix values have been z-scored across spatial bins for visualization. (G) Significance of the spatial correlations between split-session ripple fields as in (F) compared to shuffle, measured across sessions. (H) Trajectory densities for replay (top) and rat (bottom), z-scored as in (F) for visualization. (I) Significance of spatial correlations between ripple fields and replay (left) and rat (right) trajectory densities, respectively, measured across sessions. (J) Significance of the directional tuning of replays containing ripples that cross ripple field zones compared to shuffle, measured across sessions and ripple field zones (n = 45 ripple field zones; each session could contribute more than one ripple field zone – see Methods).

As can be seen in the examples of **Figure 1**, ripples tended to co-occur across tetrodes. When ripple fields when computed separately for each tetrode, they were found to be highly correlated, even across hemispheres (**Figure 3A,D**), suggesting that ripples across the hippocampus are tightly coordinated with respect to spatial content within replay. While the local LFP yielded consistent ripple fields across tetrodes, the local population spiking did not. Spatial fields derived from the local population spiking from each tetrode were highly variable across tetrodes (**Figure 3B,D**). However, the spike density field derived from all cells across tetrodes (**Figure 3C**) was strongly correlated with the ripple field (36 out of 37 sessions; **Figure 3E**), showing similar levels of spatial information and stability (37 out of 37 sessions and 33 out of 37 sessions significant, respectively). This result indicates that ripples and global population bursting are tightly yoked across replays and respond to similar spatial content within replay. It also suggests that ripples measured locally are representative of the global population dynamics of the hippocampus (Buzsáki et al., 2012) and that the inhomogeneities of the ripple field are not driven by local biases in the cell sampling.

**Figure 3.**
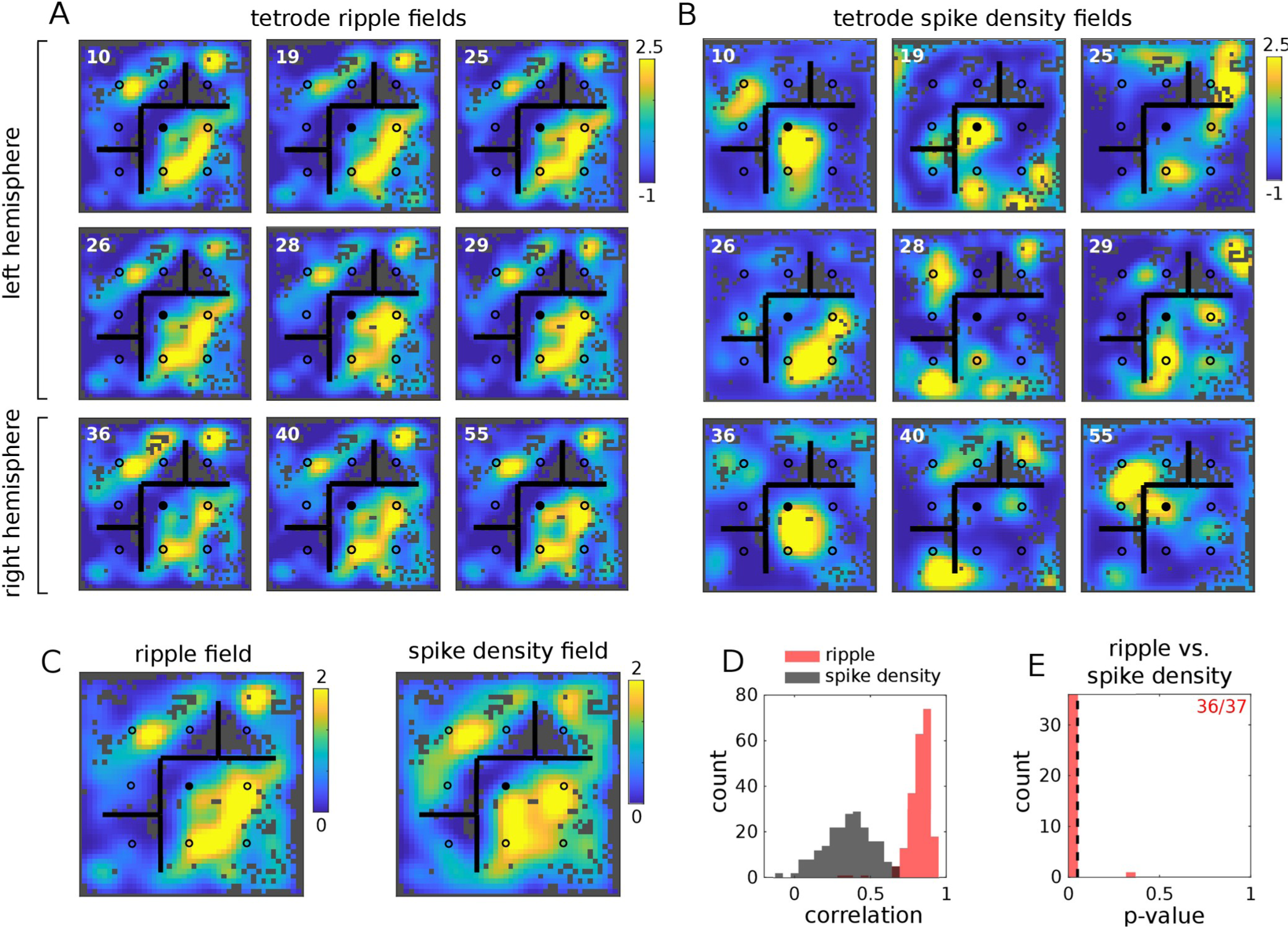
Ripple fields are replicated across tetrodes and hemispheres. (A) Ripple fields from a subset of tetrodes from the left (top two rows) and right (bottom row) hemispheres (tetrode number indicated at upper left), z-scored across matrix elements for visualization. Ripple power was computed separately for each tetrode. (B) Same as (A) except using spike density computed locally using only the cells recorded on each tetrode. (C) Full ripple (left) and spike density (right) fields computed by averaging across tetrodes or including all cells recorded across tetrodes, respectively. (D) Distribution of spatial correlations between individual tetrode and full ripple fields (red) and spike density fields (black). Tetrodes were required to have at least 5 detected place cells and mean z-scored ripple power during replay greater than 0.5 (n = 213 tetrodes). (E) Significance of correlations between ripple and spike density fields measured across sessions (n = 37 sessions). Null distribution computed by circularly permuting ripple power across replay events and measuring spatial correlations between shuffled ripple and unshuffled spike density fields. Number of significant sessions indicated at upper right.

### Ripple field locations are predicted by place field over-representation during run

Are there aspects of the environment or the animal’s behavior that correlated with the location of the ripple fields? Surprisingly, ripple fields did not correlate with the reward locations or the distance to barriers (2 out of 37 sessions and 0 out of 37 sessions, respectively; **Figure S3**). Across session pairs, ripple fields did not correlate with locations in the environment in which the local barrier structure was the same (2 out of 22 sessions) or had changed (1 out of 22 sessions), or where the animal’s behavior was similar (1 out of 22 sessions) or different (3 out of 22 sessions). Ripple fields were nevertheless highly correlated across rats for the subset of sessions in which rats experienced the same barrier configuration (7 out of 8 sessions significant; **Figure 4**), indicating that ripple field locations were nevertheless a function of the environmental structure.

**Figure 4.**
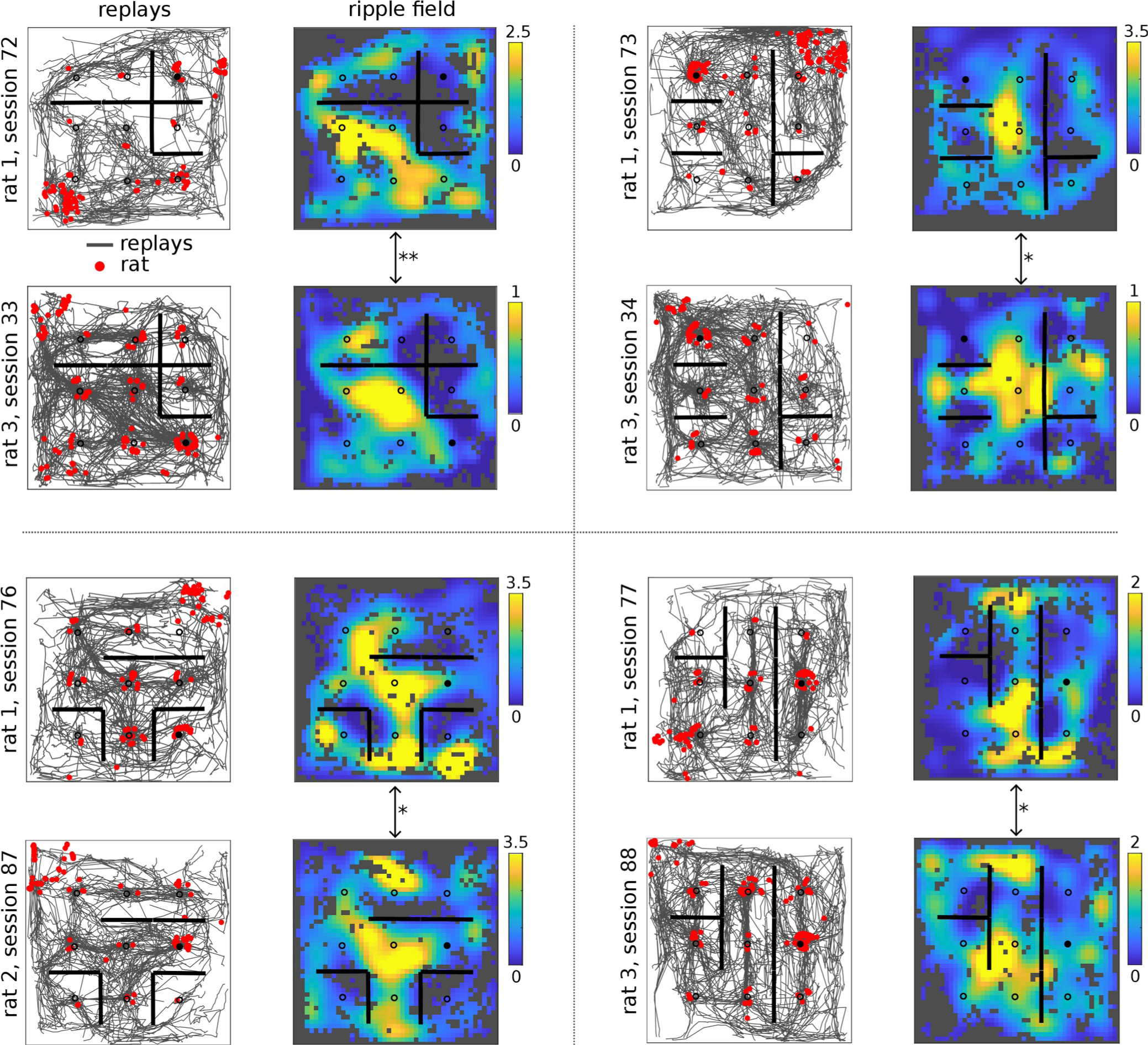
Ripple fields are similar when barrier configurations are matched across rats. Replay trajectories (left) and ripple fields (right) for pairs of rats experiencing the same barrier configuration (quadrants). Significance of the spatial correlation between ripple fields compared to shuffle is also indicated (*p<0.05; **p<0.01). Shuffles computed by circularly permuting ripple power for the second sessions across replay events and measuring spatial correlation.

We next examined whether aspects of the hippocampal representation during running predicted the location of ripple fields. Previous work has shown that during replay, individual place cells recapitulate their peak firing rates during run (Pavlides & Winson, 1989; Hirase et al., 2001; Battaglia et al., 2005; Tirole et al., 2021). To compare this at the population level, we summed the place fields during run (**Figure 5A**) and compared that distribution to spike density fields during replay (**Figure 5B**). Strikingly, these distributions were significantly correlated for every session (37 out of 37 sessions; **Figure 5D**), suggesting that cell recruitment during replay is strongly driven by the density of place fields during run. Likewise, ripple fields were less but still strongly correlated with the summed place fields (26 out of 37 sessions; **Figure 5A,C,D**). We hypothesized that the lower correlation in the ripple fields-to-place fields comparison was due to a nonlinearity in the process that converts place cell recruitment during replay into ripples, as has been suggested (Buzsaki, 2015). Indeed, peak ripple power and peak population bursting across replay events was strongly nonlinear (**Figure 5E**). Together, these results suggest that place cell over-representation during run is a strong determiner of when ripples occur during replay.

**Figure 5.**
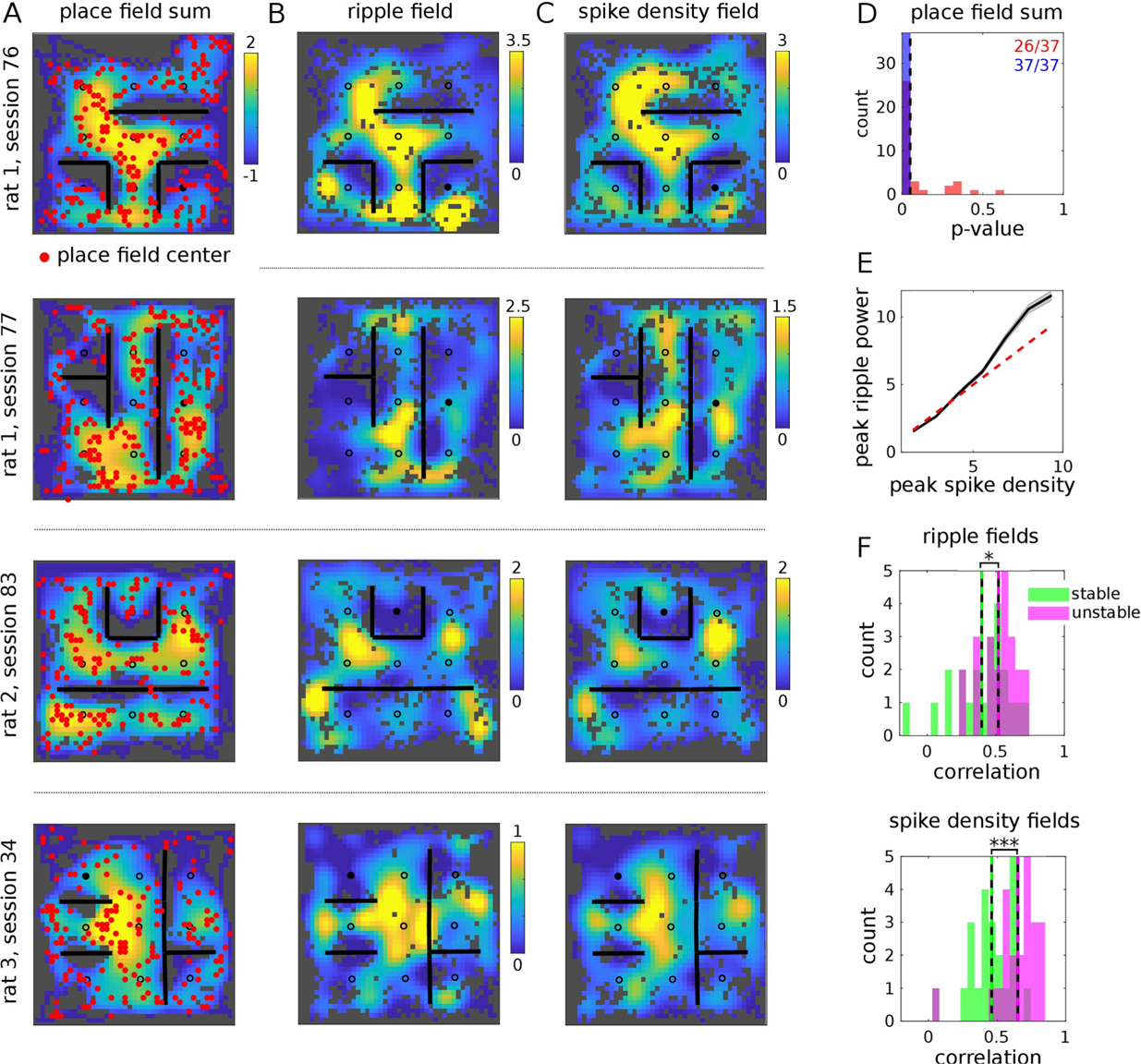
Ripple field locations are predicted by place field over-representation during run. (A) Place field rate maps summed across all cells from four sessions across three rats. Matrix values have been z-scored for visualization. Red dots indicate place field centers from all place cells. (B) Spike density fields and (C) ripple fields from the corresponding sessions in (A). (D) Significance of spatial correlations between ripple/spike density fields and summed place field maps (red/blue histograms, respectively), measured across sessions (n = 37 sessions). Null distribution computed by circularly permuting ripple power/spike density across replay events and measuring spatial correlations between shuffled ripple/spike density fields and the summed place field maps. Number of significant sessions indicated at upper right. (E) Peak ripple power vs. peak spike density across replays (n = 7273 replays). Dashed red line is the identity line. (F) Top: Spatial correlations between ripple fields and summed place field maps, for stable (green) and unstable (pink) cells. Stability is measured with respect to the same cell’s place field in previous session. Bottom: Same as top but using spike density fields (n = 22 sessions; paired t-test applied to distribution of spatial correlation values for stable and unstable cells measured across sessions; *p<0.05, ***p<0.001).

Previous work has suggested that ripples preferentially reactivate cells with remapped place fields associated with novel environments or goal locations (Cheng & Frank, 2008; Dupret et al., 2010). To probe whether ripple fields were associated with place field reorganization, we divided cells into stable and unstable groups, depending on whether the cell’s place field for the current session was significantly different from the previous session. Summed place field maps for the unstable cells were more strongly correlated with the spike density fields and ripple fields than were the summed maps of the stable cells (**Figure 5F; Figure S4**). This is striking given that unstable cells are a minority (30%) in this task (Figure 5 in Widloski & Foster, 2022). Taken together, this result shows that ripples selectively echo locations where unstable cells congregate in response to spatial manipulations of the environment.

## DISCUSSION

We have shown that replay can occur separately from ripples, and that ripples themselves are bound to the replay of specific information, in the form of certain locations and not others. Previous work has suggested that during exploration of a Y-maze, ripples tended to avoid replay of the maze choice point for replays that crossed from one arm to another, suggesting that ripple timing during replay is not completely random (Wu & Foster, 2014). Our work substantially elaborates on this finding. We were able to define ripple fields, locations encoded by replay where ripples were reliably elicited, and showed that ripple fields exhibited spatial information and within-session stability that was above chance and an independence with respect to the travel orientation of replay, thus sharing many of the essential allocentric coding properties that belong to single-cell place fields. Crucially, unlike place fields, ripples occurred independently of the rat’s location, a remarkable fact given that rat location is such a strong determiner of replay starting locations (Foster & Wilson, 2006; Diba & Buzsáki, 2007; Davidson et al., 2009). This implies that ripple occurrences are largely detached from the rat’s immediate sensory experience but nevertheless follow an internal logic that is only revealed by recording and decoding large populations of place cells in open environments.

Our results challenge the notion that ripples are needed to elicit long replays. The average duration of ripple-less replays was ∼270 msec, but we found examples of ripple-less replays that lasted well over 1 second (**Figure 1C**). We suggest that long, ripple-less replays exist by virtue of the fact that they avoid traveling into ripple zones. On linear tracks, traveling through ripple zones may be unavoidable, which is why replays are nearly always accompanied by ripples (see Figure 5A in Davidson et al., 2009) and why replay duration is strongly correlated with ripple occurrence (Davidson et al., 2009; Yamamoto & Tonegawah, 2017). While the latter was also true in our data set, this is simply explained by the fact that long replays have, by chance, increased likelihood to cross through multiple ripple zones.

The large prevalence of ripple-less replays, which make up about 65% of all replays according to our criteria, has two further implications. First, it suggests that studies investigating causal manipulations of replay through the detection of ripples (Jadhav et al., 2012; Singer et al., 2013; Girardeau & Zugaro, 2009; Maingret et al., 2016; Ego-Stengel & Wilson, 2010; Aleman-Zapata et al., 2022), especially those performed in more open environments, could be missing a great deal of sequential replay-like content, which could lead to underestimates of the effects of replay disruption on behavior. Second, it challenges the assumption that every offline sequenced reactivation necessarily participates in brain-wide systems-level consolidation. Instead, our results suggest a more hierarchical view of sequence-based retrieval processes operating in the hippocampus, with ripple-less replays being the default mode of offline hippocampal reactivation and used for possibly maintaining the local hippocampal map (e.g., Babichev et al., 2019) and its connections to cortex (Kali & Abbott, 2004), and with replays coincident with ripples participating in the integration of memory traces associated with environmental novelty and change into distributed cortical networks.

An interesting consequence of our finding is that, to the extent that ripple expression is gated by individual spatial fields, the patterns of neural activity driven by ripples may likewise resemble relatively static representations rather than sequential trajectories. This is consistent with the finding that individual neurons in the prefrontal cortex exhibit activity during hippocampal ripples that is selective for which arm of a Y-maze is being replayed but not for individual locations within the arm (Berners-Lee et al., 2021). Likewise, VTA neurons have been shown to selectively fire at replayed reward locations during ripples (Gomperts et al., 2015), while both the retrosplenial and secondary motor cortices show selective activation to landmarks and cues during reactivations during rest (Chang et al., 2020; Chang et al., 2023). Taken together, these results suggest that what is broadcast from the hippocampus during ripples is a more narrowly restricted set of experiential content than replay, which could explain selective memory for salient experiences (LaBar & Phelps, 1998; Kensinger, 2009).

Our unique behavioral paradigm using repeated spatial and reward manipulations allowed us to probe the relationship between ripple fields and behavioral context. While ripple field locations were replicated across animals experiencing the same context, they were not tied to reward locations coinciding with rewards or manipulated barriers. Instead, ripple fields were strongly correlated with the density of place fields observed during run, especially locations where the unstable cells congregated in response to the barrier and reward manipulations. This supports the idea that ripples specialize in broadcasting information related to environmental change and is consistent with previous work showing that ripples promote the stabilization of new spatial representations in the hippocampus associated with novel environments, novel reward contingencies, or novel goal locations (Cheng & Frank, 2008; Singer & Frank, 2009; Dupret et al., 2010; Hwaun & Colgin, 2019; Roux et al., 2017; Csicsvari & Dupret, 2014). More broadly, the tight relationship between ripple fields and place field overrepresentation suggest a novel role for the nonlinear reverberatory process thought to underlie ripple generation (Buzsáki, 2015), as reflected in the nonlinear relationship between population spiking and ripple power during replay that we measure. This mechanism may create the spatial distribution of ripple fields as a direct result of where in the environment place fields are over-represented.

Together, our results dissociate two experimental phenomena, ripples and replay, long considered to be aspects of the same process. Moreover, they suggest a novel component of hippocampal-dependent systems memory consolidation, by which certain experiences and not others are selected for further processing. Understanding the functional significance of this relationship will be crucial for elucidating the full spectrum of cognitive processes supported by hippocampal replay.

## METHODS

Neural activity was recorded from dorsal hippocampus (region CA1) of 3 Long-Evans rats (*Rattus norvegicus*; 3-4 months old) performing a goal-directed task in an open field maze with movable barriers (task described below). Rats were housed in a humidity and temperature controlled facility with a 12-hour light-dark cycle. Before the start of the experiments, rats from the same breeding cohort were housed in pairs. At the start of the experiments, rats were single-housed. All experimental procedures were in accordance with the University of California Berkeley Animal Care and Use Committee and US National Institutes of Health guidelines.

### Task design and training

Rats were trained on a spatial memory task in a square arena to search for liquid chocolate available in one of 9 food wells, which alternated on consecutive trials between a learnable fixed location (“Home” well) and unpredictable other locations (“Random” wells), designated as Home and Random trials respectively. On each trial a variable time delay (5-15 sec) passed before: (a) reward was provided at the bait location, and (b) for all Random trials, a light came on next to the rewarded well, cueing the approach. Before each session, transparent “jail-bar” barriers (Olafsdottir et al., 2015), permeable to visual and olfactory information, were placed in 6 out of 12 possible locations, in a novel, random selection from 924 possible configurations. 2-3 consecutive behavioral sessions were performed per day, each separated by ∼3-4 hours, and each with a novel barrier configuration (or in some cases, no barriers) as well as a novel, pseudo-randomly chosen Home location (47 sessions total; 12 sessions for rat 1, 17 sessions for rat 2, 8 sessions for rat 3).

### Drive design and surgery

Rats were implanted with microdrive arrays weighing 40-50 grams and consisting of 64 independent-adjustable tetrodes made of twisted platinum iridium wires (Neuralynx) gold plated to an impedance of 150-300 MOhms. Drive cannulae were implanted bilaterally to target hippocampal dorsal CA1 (−4.13 AP, 2.68 ML relative to bregma) using a surgery protocol described elsewhere (Pfeiffer and Foster, 2013; Widloski and Foster, 2022). Tetrodes were slowly lowered to the cell layer over the course of 2-4 weeks, which was identified by the presence (and shape) of strong-amplitude sharp-wave ripples. The rats were allowed 3-4 days recovery, after which behavioral training on the barrier maze task was resumed but without food restriction in their home cages. Food restriction was resumed a week after surgery.

### Behavioral analysis

Rat position was tracked using automated software from Spike Gadgets and sampled at 30 Hz. Position and velocity were smoothed using a Butterworth filter (second order with a cutoff frequency of 0.1 samples/s using the butter function in Matlab, selected to give reasonable smoothing to the rat’s trajectory). After smoothing, positions were interpolated at 200 Hz (temporal step size of 5 msec) so as to match the sample rate of decoded points within replay (see section Replay detection below)

### Cluster analysis

Spikes were extracted from channel LFP’s (sampled at 30 kHz) using Spike Gadgets Trodes software and clustered automatically using Mountainsort (Chung et al., 2017) and merged across sessions using the *msdrift* package. Additional cluster mergings across sessions was performed manually based on similarity of waveform. Clusters were accepted if noise overlap < 0.03, isolation > 0.95, peak SNR > 1.5 (Chung et al., 2017) and had passed a visual inspection.

### Place fields

In order to compute the cell’s *place field*, for each spike that occurred during run (rat speed > 10 cm/s), the rat position was found through linear interpolation (*interp1* in Matlab). Positions were binned with 2 cm square bins. The unsmoothed rate map for the *i^th^* cell was defined as

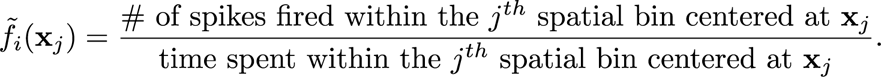

Smoothed rate maps, denoted as *f_i_*(**x***_j_*), were computed by first setting unvisited bins to zero and convolving the rate maps with a 2D isotropic Gaussian kernel (8 cm standard deviation (SD)).

Spatial information (bits/spike) for the *i^th^* cell was defined as

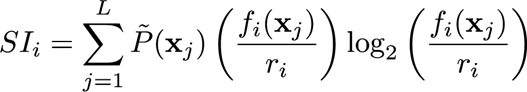

where L is the number of spatial bins, 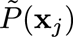 is the probability of the rat or replay being at the *j^th^* spatial bin, and 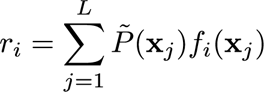 is the cell’s mean firing rate. *Place cells* were identified as having place fields with *r* > 0.01 Hz and *SI* > 0.5 bits/sec.

To determine place field stability, first rate map correlations were defined as the Pearson’s correlation between any pair of rate maps (i.e., place fields). For the *i^th^* cell, with rate maps 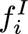 and 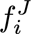 for the *I^th^* and *J^th^* sessions respectively, the rate map correlation was

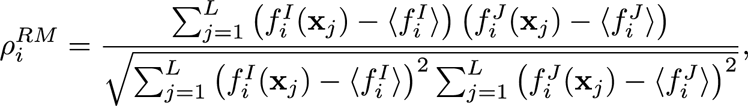

where 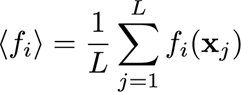 is the mean spatial firing rate. Rate map correlations were evaluated only at visited spatial bins common to both sessions and were only measured for cells identified as place cells for both sessions and whose rate maps had minimal barrier overlap in both sessions. Barrier overlap was assessed as follows: First, spatial bins were denoted as “active” if the firing rate density in that bin was greater than 1 Hz/cm. Rate maps for which at least 60% of all active bins were at least 4 cm away from the nearest barrier were considered to minimally overlap with the barriers. A rate map correlation shuffle distribution was computed for each place cell by randomly permuting place cell ID’s 100 times in the second session and recomputing the correlations. A place cell was called stable across a pair of sessions if its rate map correlation exceeded the 95th percentile of its shuffle distribution; otherwise it was called unstable. Field centers were calculated using a density-based clustering approach. First, rate maps were treated as discrete probability distributions and resampled 2500 times (using the *pinky* function in Matlab). Then, the sample points were clustered using *dbscan* in Matlab, with a neighborhood search radius of 2.5 bins and a minimum number of neighbors of 50. Field centers were calculated as the center-of-mass of all points belonging to the same cluster.

### Bayesian decoding

Let *k_i_* be the number of spikes emitted by the *i^th^* place cell in a given time bin of duration τ. The posterior probability at bin **x***_j_* conditioned on the activity vector 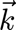 (with the *i^th^*element as *k_i_*) is given by Bayes rule (assuming Poisson spiking noise statistics, independence between neurons, and a uniform spatial prior (Davidson et al., 2009)):

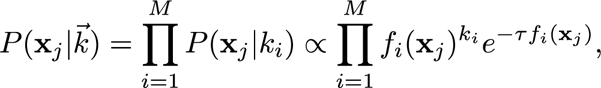

where *M* is the number of neurons. A uniform prior was used for the purposes of making minimal assumptions about the location of the decoded positions. The posterior probability was computed for all bin locations **x***_j_* where 1 ≤ *j* ≤ *L* and *L* is the total number of spatial bins. Define 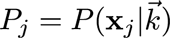 and let **x***_j_* = (*x_j_*, *y_j_*) be the components of the *j^th^* spatial bin. The components of the *posterior center-of-mass* (COM) were given by

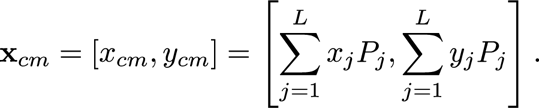

The *posterior spread* was defined as the square root of the second central image moment of the posterior:

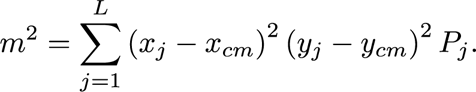

The *posterior COM jump size* was defined as the L2 norm of the difference vector between consecutive posterior center-of-mass estimates:

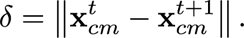

### Replay detection

We briefly outline the procedure for detecting replays, which has been previously published (Widloski and Foster, 2022). The Bayesian decoder was applied to spikes within a sliding window of 80 msec duration (shifted in 5 msec increments) over the entire session from all place cells found in the session. Time bins were kept for further analysis based on three criteria: rat speed (*v_rat_* < 5 cm/s; rat speed was computed at the center of each time bin via linear interpolation), posterior spread (*m* < 10 cm), and posterior COM jumps size (δ < 20 cm). We defined a *sub-sequence* as a set of temporally contiguous bins satisfying the above criteria. Sub-sequences captured epochs in which the posterior was well defined (small posterior spread) and moved smoothly (small COM jump size across time steps). Neighboring sub-sequences were merged if the spatial and temporal gap between them was 20 cm and 50 msec, respectively. A sub-sequence (merged or not) was denoted a *candidate sequence* if its duration was greater than 100 msec. *Replays* were selected as candidate events that were sufficiently spatially dispersed. To this end, we defined a *spatial dispersion* metric:

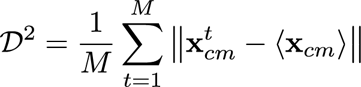

where *M* is the length of the sequence and 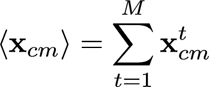. A candidate sequence was defined as a replay if its dispersion was greater than 12 cm.

### Ripple amplitude and spike density

Sharp wave-ripple amplitude was computed for each tetrode by band-pass filtering the LFP in the 120 to 170 Hz range, extracting the amplitude envelope via a Hilbert transform, and then averaging the amplitude across tetrodes. Population spike density was computed by first summing the total number of spikes from all clusters in 1 msec non-overlapping time bins. Both the ripple amplitude and spike density were smoothed through convolution with a Gaussian kernel (80 msec SD) and z-scored. The mean and standard deviations used for z-scoring were computed from stopping periods only (i.e., rat speed < 5 cm/s).

Ripple events were detected as when the z-scored ripple power peaked above 3 standard deviations. The time boundaries of each ripple event were determined by the nearest dip in the ripple power before and after the ripple peak time. Adjacent ripple events were merged if the time boundaries were less than 50 msec apart. Ripple events were required to have a duration of at least 50 msec. A given replay was determined to be ripple-less if it (1) contained no ripple events (according to the ripple detection procedure) and (2) passed a ripple power shuffle test requiring that the peak ripple power within the replay exceed the 95^th^ percentile of a shuffle distribution comprised of ripple power peaks taken from 100 random equal-length snippets of LFP during stopping periods but outside of ripple times.

### Ripple and spike density fields

The mean ripple amplitude and spike density as a function of replay location (i.e., “ripple field” and “spike density field” respectively) were computed across spatial bins (2 cm bins, same as for place fields) and normalized by replay occupancy. As with place fields, smoothed rate maps were computed by first setting unvisited bins to zero and convolving the rate maps with a 2D isotropic Gaussian kernel (8 cm standard deviation (SD)). *Ripple field zones* were calculated using the same density-based clustering approach used to find place field centers. First, ripple fields were treated as discrete probability distributions and resampled 10,000 times. Then, the sample points were clustered using *dbscan* in Matlab, with a neighborhood search radius of 3 bins and a minimum number of neighbors of 250. Only clusters with at least 1,000 samples were considered. For each of these clusters, the boundary was taken as the level set of the cluster density equal to 0.3 of the peak value. After defining the ripple zone boundary, replay passes through the boundary were extracted. Only passes with a minimum length of 30 cm were considered. The direction associated with each replay pass was computed as the average angle of all segments of the pass (replays were sampled at intervals of 5 msec as described above).

### Local barrier similarity

Local barrier similarity is a measure of the local environmental structure across two barrier configurations, as defined in Widloski & Foster, 2022. First, the *barrier potential* was computed for each barrier configuration by convolving the barriers with a 2D isotropic Gaussian kernel (10 cm SD). Let 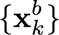 be the set of bin overlapping with the barriers. The barrier potential at the *i^th^* spatial bin was computed as

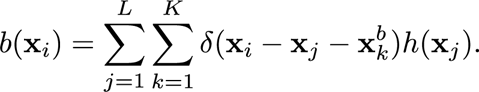

where *h* is the Gaussian kernel and *K* is the number of barrier-overlapping spatial bins. We defined the *Local Barrier Similarity* (LBS) at the *i^th^* spatial bin across a pair of sessions I and J as

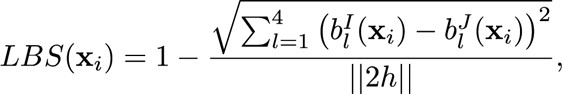

where the summation is over the 4 closest spatial bins to the *i^th^* bin and ||.|| is the L2 norm.

### Local rat trajectory similarity

Like the local barrier similarity, the local rat trajectory similarity measures the local similarity of the rats trajectory across two barrier configurations. First, for each session and for each spatial bin, the distribution of angles for all segments of the rat’s trajectory that exist inside a circle of radius 8 cm centered around that bin were computed. The local rat trajectory similarity across a pair of sessions was computed as the correlation between the two angle distributions at each spatial bin.

## QUANTIFICATION AND STATISTICAL ANALYSIS

Statistical tests and corresponding p-values are reported within the figure legends. All statistical analyses were performed in Matlab. Paired-sample t-tests were used for paired comparisons and two-sample t-tests for non-paired comparisons. Binomial tests were performed assuming 50% chance occurrences.

## DATA AND CODE AVAILABILITY

All data and custom-written code are available upon request.

## ACKNOWLEDGEMENTS

We would like to thank Matt Kleinman and Caitlin Mallory for helpful discussions.

## FUNDING

This work was supported by NIH grant NS113557. Animal use conformed to NIH guidelines and was approved by the UC Berkeley Animal Care and Use Committee.

## AUTHOR CONTRIBUTIONS

JW and DJF conceived of and designed the study. JW acquired the data and performed the analyses. JW and DJF wrote the manuscript.

## DECLARATION OF INTERESTS

Authors declare that they have no competing interests.

## SUPPLEMENTAL INFORMATION

**Figure S1, in relation to Figure 1.**
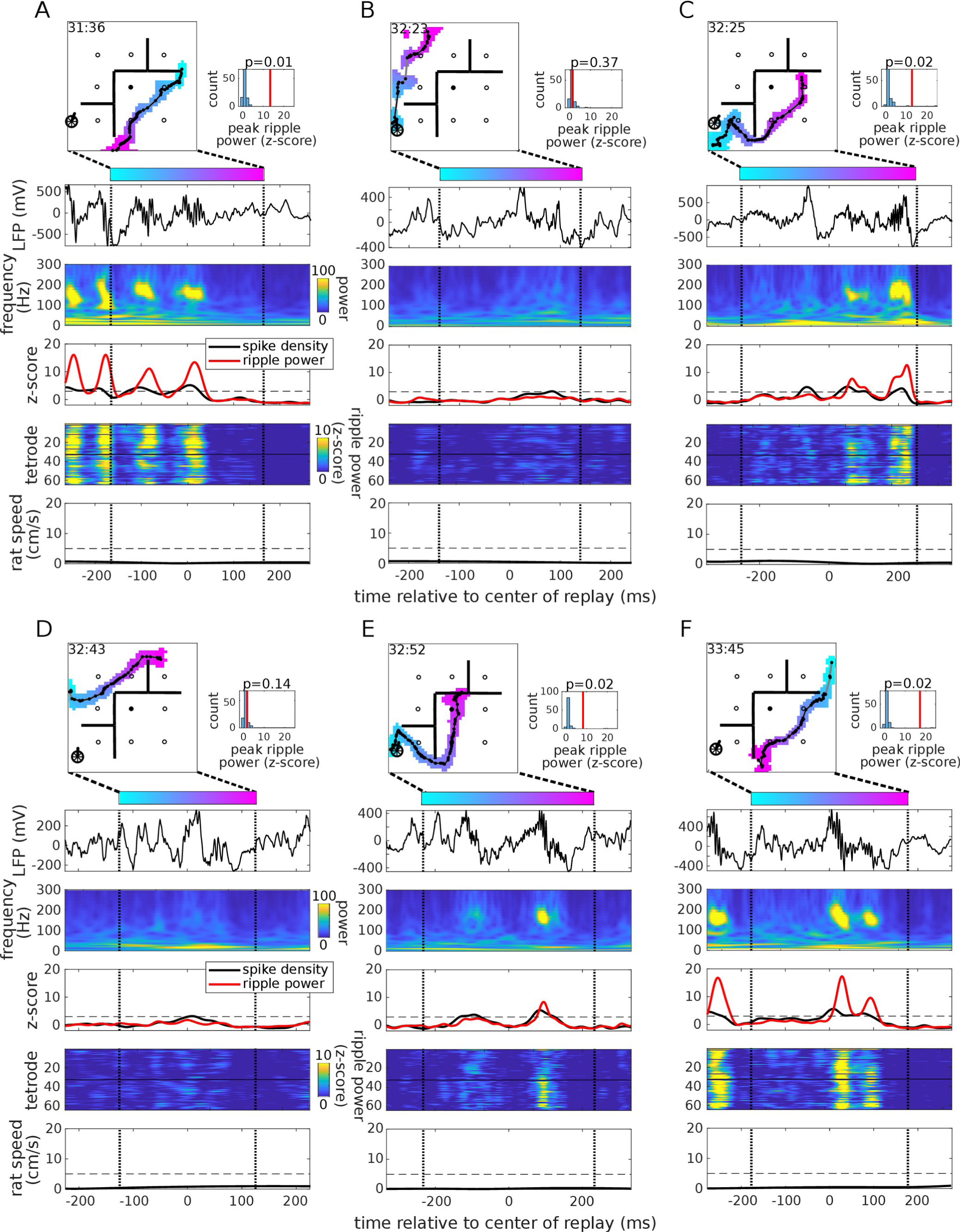
(A) Top row: Example replay from rat 1, session 75. The colored blob is the posterior probability of a replay thresholded at 0.01 and color-coded according to elapsed time within the replay. Solid black line: replay center-of-mass. Time within session is shown at the upper left (min:sec). Black circles and straight lines are the locations of reward and transparent barriers, respectively. Filled circle is the home well. Right: Peak ripple power (vertical line) vs. shuffle, where shuffle is the peak ripple power (z-scored and averaged across tetrodes) from 100 random equal-length snippets of LFP during stopping periods but outside of ripple times, with p-value indicated above. Second row: Raw LFP trace from single electrode as a function of time within the replay. Black vertical dashed lines demarcate start and end of replay. Third row: Spike density summed across all recorded cells (black) and ripple power averaged across tetrodes (red). Dashed horizontal line indicates z-score of 3 and is standard detection criteria for ripple/spike density events. Fourth row: Ripple power across tetrodes. Tetrodes 1-32 (33-64) targeted the left (right) hemisphere. Fifth row: Rat speed. Dashed horizontal line indicates rat speed of 5 cm/s. (B-F) Replays recorded during the same stopping period.

**Figure S2, in relation to Figure 2.**
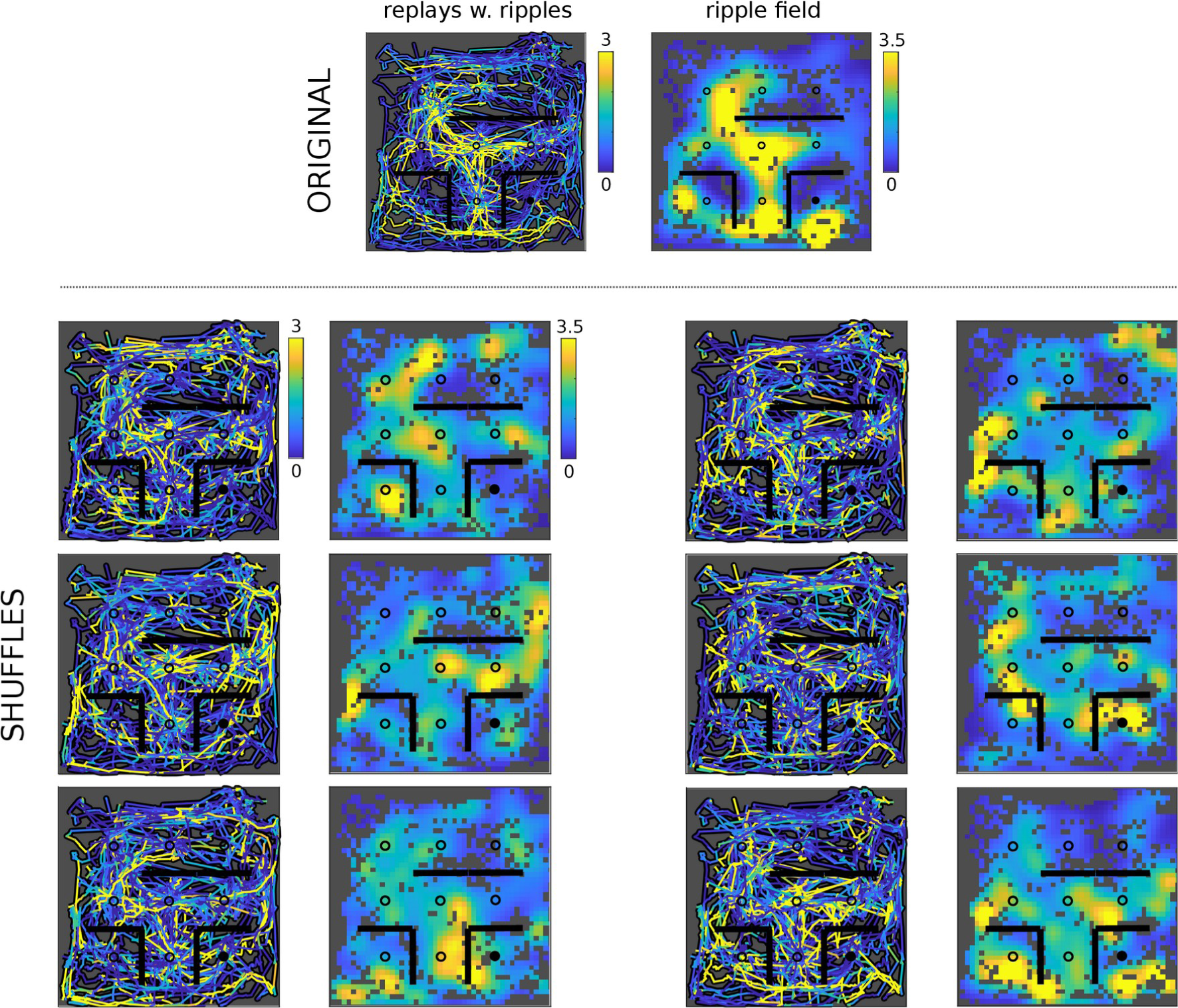
Top: Replays with ripple power superimposed (left) and corresponding ripple field (right) for session 76, rat 1. Bottom: 6 shuffle examples, where for each pair, ripple power has been circularly permuted across all replay events. Color scale bars are the same for all shuffle examples.

**Figure S3, in relation to Figure 4.**
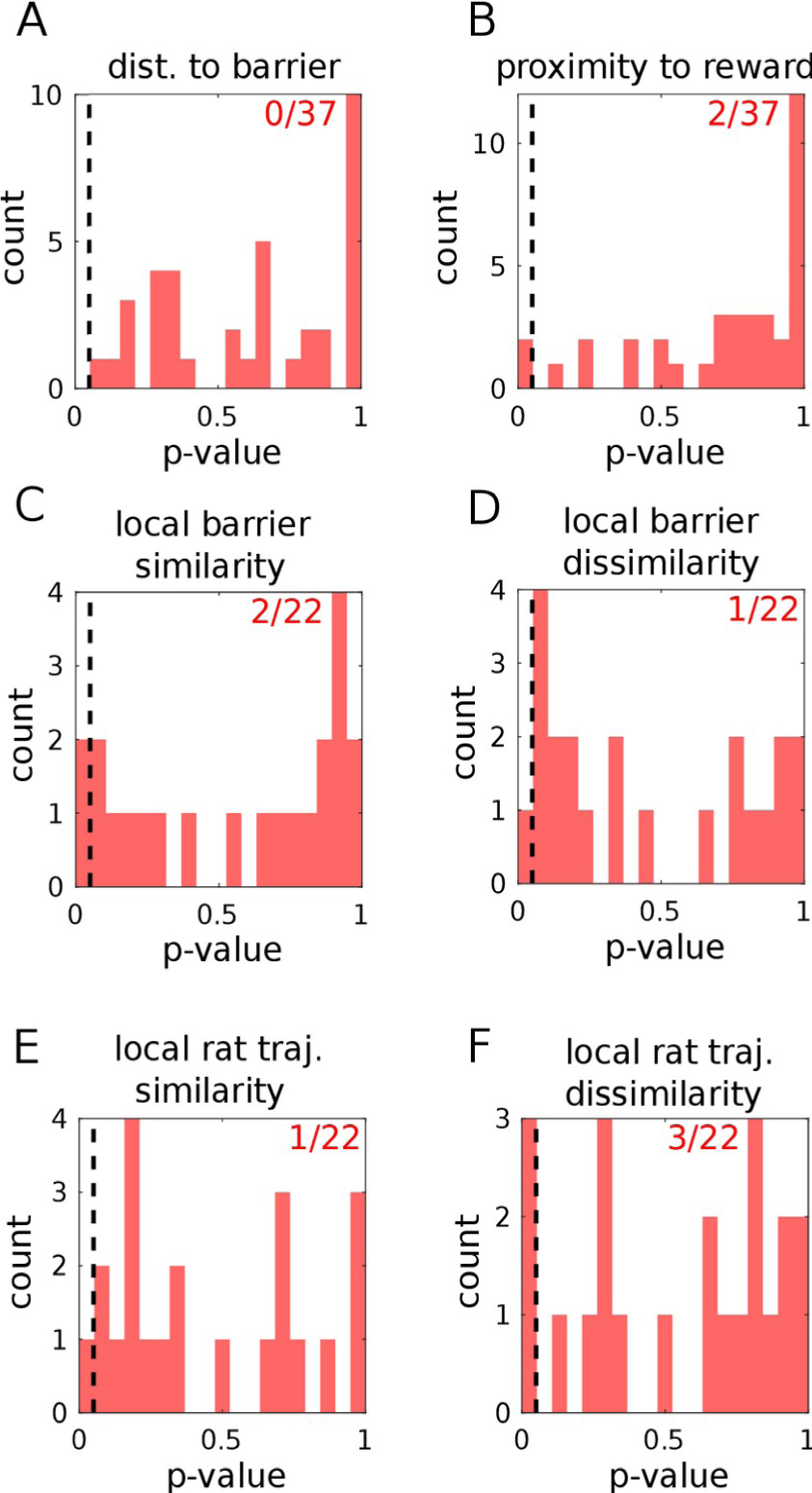
Significance of ripple field correlations with distance to the nearest barrier (A), proximity to nearest reward locations (B), local barrier similarity between the current and previous session (C), local barrier dissimilarity (D), local rat trajectory similarity between the current and previous session (E), and the local rat trajectory dissimilarity (F), measured across sessions (n = 37 sessions).

**Figure S4, in relation to Figure 5.**
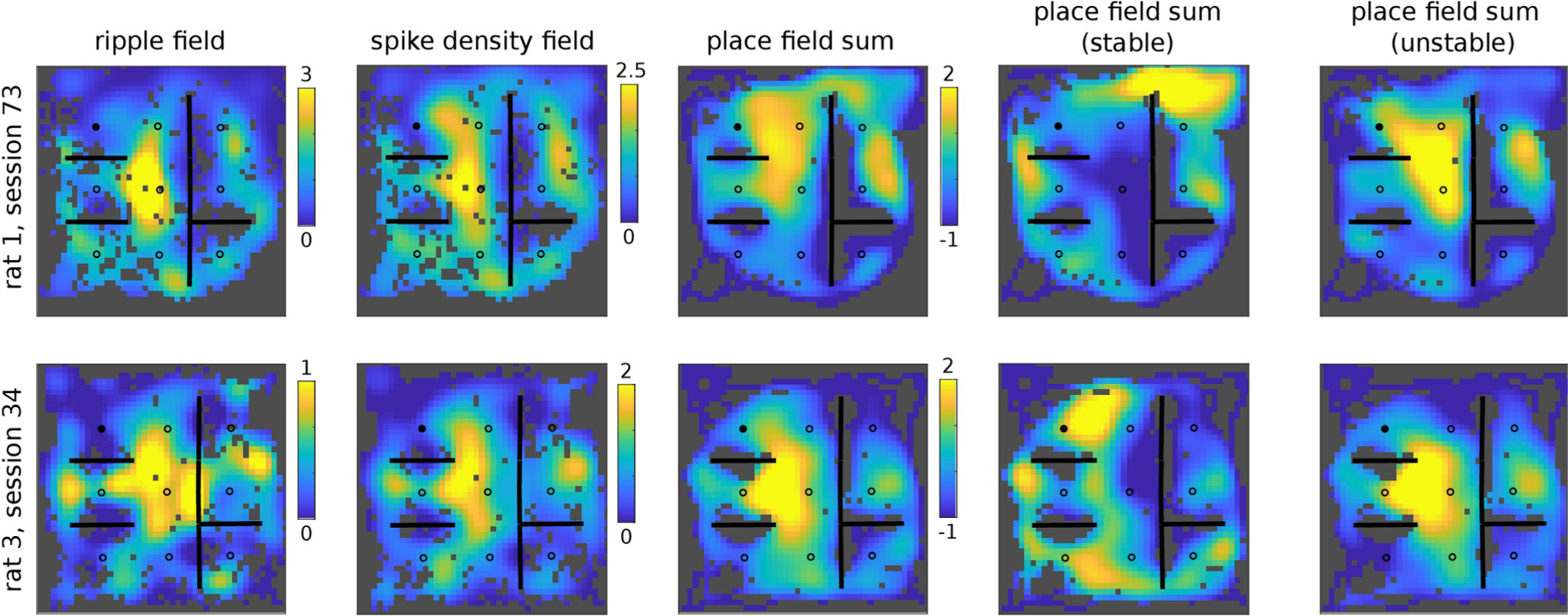
Comparison of ripple fields with summed stable and unstable place fields, for two rats experiencing the same session. From left to right: Ripple field, spike density field, place field sum, and place field sum for stable and unstable cells. The matrix values for the last three columns (place field sums) have been z-scored for visualization and have the same color axis limits. Top row: rat 1, session 73; Bottom row: rat 2, session 34.

## Notes

### Competing Interest Statement

The authors have declared no competing interest.

